# Speech-like sounds dominate the human infant vocal landscape

**DOI:** 10.1101/2021.01.08.425949

**Authors:** D. Kimbrough Oller, Gordon Ramsay, Edina Bene, Helen L. Long, Ulrike Griebel

## Abstract

Early human infant vocalization is viewed as forming not only a critical foundation for vocal learning of language, but also a crucial realm of communication affecting emotional and social development. Although speech-like sounds are rare or absent in other ape infants, they share distress sounds (shrieks and cries) and laughter with humans, forming a potential basis for especially informative cross-species comparisons as well as potential insights regarding usage and learning of vocal sounds. A fundamental need to make such comparisons possible is empirical research to document frequency of occurrence of vocalizations of various types in natural environments.

The present work focuses on laughter in the human infant, a topic that has been viewed by many as a key factor in social development for humans and other apes. Yet we know of no research quantifying frequency of occurrence of human infant laughter in natural environments across the first year. In the past two decades it has been shown that the predominant vocalizations of the human infant are “protophones”, the precursor sounds to speech. Longitudinal research has indicated unambiguously that protophones outnumber cries by a factor of at least five based on data from random-sampling of all-day recordings across the whole first year. The present work expands on the prior reports by reporting data showing that human infant laughter occurs even more rarely than cry in all-day recordings. Yet laughter is clearly a salient and important aspect of social development. We reason about the dominance of protophones in the infant vocal landscape in light of their role in illuminating human vocal learning and the origin of language.

## Background and rationale

The pursuit of the roots of vocal learning in various taxa is hoped to provide perspective on how human vocal learning evolved and eventually provided the basis for language. Yet surprisingly little is known about many issues where empirical grounding is required in order even to begin the process of insightful cross-species comparisons. A variety of surprises have emerged through recent research about human infant vocal development. These surprises suggest a need for reorienting our views about vocal development and vocal learning in humans. Also, the results suggest need for new approaches to quantitative comparative research on possible foundations of language as seen in other animals [1].

There has been considerable emphasis in work on language development on a role for acquisition by copying, with caregiver interaction and modeling driving imitation [2, 3], a process whereby infants are presumed to absorb the native language. Vocal imitation is thought to begin at birth [4], and the emphasis on parental “input” and infant decoding and copying would seem to supply, in this viewpoint, a primary method by which language is learned [5]. Considerable research supports a theory of language acquisition where the focus is on perceptual learning from the language environment facilitated by parental speech [6, 7]. The literature on infant-directed speech and its potential role in language acquisition is massive [8-14]. A contrasting view of course proposes that language learning requires human-specific genetic underpinnings and downplays the possibility that language input is structured in such a way as to allow learning primarily by copying [15]. This view, however, has been heavily critiqued, and a “constructivist” point of view has been proposed as an alternative [16-19].

Ethological research, some of it quite recent, suggests a view that does not highlight imitation or vocal responsivity to parental input, but rather emphasizes human infant endogenous exploratory activity as a primary driver of very early development of language foundations. The infant’s exploratory activity, observable from the first day of life, appears to contradict several widespread expectations about how the foundations of language emerge. One of the most salient recent findings is that human infant vocal development does not begin with an overwhelming predominance of cry. On the contrary, longitudinal ethological research involving both laboratory and home recordings has illustrated that even in the first month of life and throughout the first year, cries occur far less frequently than the precursors to speech, the “protophones” [20, 21]. Additional empirical evidence on this point will be presented in the present work. Yet cry has been thought by some to be the foundation for speech, the basis upon which speech sounds develop [22]. Even in infants born prematurely by more than two months, still in neonatal intensive care, protophones outnumber cries substantially, from shortly after de-intubation, thus as soon as the infants are able to breathe on their own [23].

The sheer number of protophones produced by the human infant may seem astounding, having been determined by coding of randomly-selected samples of all-day recordings to be ∼3500 per day, a number that varies little across the first year [24]. The rate of protophone production is not discernibly lower when infants are alone in a room than when they are engaged in protoconversation. Perhaps even more surprisingly the great majority of protophones are not directed toward any listener even in circumstances where caregivers are talking to babies [25].

While vocal imitation is a logically necessary capacity for learning a lexicon, it is rare that infants actually produce immediate vocal imitation in the first year as observed in longitudinal observation [26-28]. Attempts to experimentally show such imitation are fraught with ambiguities of interpretation as to whether the infant imitates or the parent induces and/or follows the infant’s vocal explorations, a kind of following that can yield a false impression of representational imitation on the part of the infant. A systematic attempt to identify auditorily recognizable cases of infant vocal imitation for research on listener judgments of degree of imitativeness yielded no more than a handful of clear cases out of over 6000 utterances drawn from recordings of mother-infant interaction, with fewer than 5% showing *any* discernible imitation [29]. The cases that did show some discernible imitation included apparent matching of subtle prosodic features subject to notable coder disagreement regarding imitativeness. Further the mother’s presumed model utterances might have constituted productions by her of sounds she knew to be in the infant’s spontaneous repertoire, sounds that were likely to be produced by the infant with or without the mother’s model.

These facts support an upgrading of our view of early vocal and language learning to envision infants as creators more than copiers. An additional finding that suggests endogenous activity more than copying of acoustic models as driving infant vocal learning is seen in the vocalizations of the congenitally profoundly deaf. Again contradicting widespread speculation, longitudinal research has shown that deaf babies produce protophones at rates that are comparable to (if not a little higher than) the rates in hearing infants [30-33]. Furthermore the types of protophones produced prior to the onset of canonical babbling (CB, exemplified by baba, dada, and so on), appear to include the whole range of types (squeals, growls, raspberries, vocants and so on) seen in hearing infants [34]. The late onset of CB in deaf infants [35-37] does not necessarily suggest that CB is learned by imitation—we see no way to rule out the possibility that CB emerges as a self-organized product of prior protophone exploration, a reflection of infant learning of vocal and articulatory patterns through auditory and/or kinesthetic awareness.

The present paper adds empirical data on a case where another human vocal act, thought to play a key role in social adaptation, occurs far less frequently than one might expect in the first year of life, as compared with the massive frequency of protophone exploration. The topic is laughter, which we and many others recognize as providing a fundamental basis for human vocal interaction [38-42]. Based on three sources of longitudinal data on early vocal development, we compare rates of laughter (which begins in human infants around 4 months of age) with rates of both protophone production and crying.

## Methods

### The Atlanta data source

As part of a consortium effort to compare infant and young child development in infants at risk and not at risk for autism, Emory University and the Marcus Autism Center in Atlanta, GA have for years been acquiring all-day recordings using the LENA battery-powered device [43, 44] from infants across the first year of life. Mothers and infants were recruited through methods that have been described extensively a prior publication’s Supplementary Materials [45]. Participation was always dependent on written informed consent from the parents in accord with permission from the Emory University IRB.

For the present study, we shall focus on 53 of those infants, for each of whom an average of 8.9 all-day recordings were obtained across the first year. All these infants have been confirmed to have “no clinical features” (no developmental disabilities) based on evaluations at 36 months. Human coding has produced data on rates of production of the three groupings of infant sounds to be reported for each recording (protophones, cry/whimper, and laughter). Human coding was conducted in Memphis in an ongoing collaboration between the institutions with IRB permission from both Emory University and the University of Memphis.

### The Memphis data source

In a separate effort, intense longitudinal research on 12 human infants has been conducted in Memphis over the past 10 years. Again, recruitment was conducted for pregnant women in accord with approval from the University of Memphis IRB, and written informed consent was provided by all the parents. Typical development was confirmed using developmental milestone questionnaires. The Memphis research has produced two kinds of data relevant to the present report: First, each infant was recorded in a laboratory setting across the first year, and second, each infant was recorded using the same LENA all-day recording method as in Atlanta. For each of the 12 infants, both laboratory and all-day home recordings yielded data at six age points across the first year. Again, human coding in Memphis provided the data on the three groupings of infant sounds.

### All-day recordings

The battery-powered LENA recorder can be placed in the vest pocket of infant clothing to produce up to 16 hours of continuous audio at a 16 kHz sampling rate. The microphone is nominally 5-10 cm from the infant’s mouth, offering high signal to noise ratio for the infant voice under most circumstances.

The device has been utilized in many thousands of recordings since 2007-8, when it first became available [46]. It has generated a new perspective on vocal development and caregiver-infant interaction by opening the door to truly representative sampling. The new perspectives include, for example, revelation of notable differences between the patterns of caregiver-infant vocalization observed in standard laboratory or other short-term recordings as opposed to in the all-day recordings offered by the LENA system [24]. Importantly, parents have been shown to produce several times more infant-directed speech in the standard recording situations than they do in the presence of wakeful infants in randomly-sampled segments from all-day recordings in the home.

Many have used the automated analysis system offered by the LENA Foundation’s software package in research on vocal development [47-49], but the work reported here is based on the more labor-intensive method of human coding of randomly sampled five-minute segments across each recording. Human coding is the gold standard for development of automated analysis of vocalizations, and in any case, two of the issues at stake in the present work, the rates of infant laughter and rates of infant-directed speech (IDS), are not counted directly by the LENA automated system.

In both Atlanta and Memphis, parents placed a fully-charged and activated recorder in a vest worn by the infant at wake-up time and left it running until bed time. During naps or bath time, the recorder was removed from the vest and left running in a location as near the infant as practical and was then placed back in the vest afterwards. The instructions encouraged parents to record in the home with no changes in the normal pattern of interaction and caregiving. The precise procedures for recording are described in detail in prior publications [23, 45].

### Laboratory recording method

The 12 infants in the Memphis study were also recorded across the first year in a laboratory designed to resemble a child’s playroom. There were eight cameras, one placed high and one placed low in each corner of the room. High fidelity wireless microphones were worn in an infant vest and at the parent’s lapel, recording at 48 kHz, with video subsequently synchronized with frame-level accuracy to the high-fidelity audio from the two microphone channels. Two channels of video (from among the 8 cameras) were selected at each point in time by staff in the adjacent control room, providing one view of the infant face and another of the interaction.

The laboratory recordings were typically one-hour in duration although sometimes the sessions were broken up into smaller segments with temporary interruptions to accommodate feeding or infant discomforts. Scheduling avoided times when an infant would be likely to fall asleep, but on occasion, especially with the youngest ages, sleep also interrupted recordings. The protocol for recording involved three segments of nominally 20 minutes duration each. These were roughly counterbalanced in order of occurrence. In one circumstance (No Adult Talk) the parent was seated in the room, reading or engaging in some other silent activity while the infant was nearby, often playing. In another circumstance (Adult to Adult Talk), the infant was nearby in the room, while the parent was engaged in a verbal interview with a staff member of the research project. In a final circumstance (Parent Infant Talk), parent and infant interacted playfully, with considerable IDS.

The laboratory recordings at the same six ages as for the LENA recordings were human coded in Memphis according to the procedures described below.

### Sample selection for coding

21 and 24 randomly-selected five-minute segments were extracted from each all-day recording from Atlanta and Memphis respectively. These segments were subject to human coding as specified below. After coding, some segments were excluded from analyses because the infant was deemed to be asleep by the coders, yielding 7387 five-minute segments from the 474 all-day recordings of the 53 Atlanta infants and 1185 from the 69 all-day recordings of the 12 Memphis infants. The 67 human-coded Memphis laboratory recordings were approximately one-hour each: all 12 infants had recordings at five of the six ages across the first year, but only 7 had recordings at the youngest age. The laboratory recordings yielded 59, 66, and 64 sessions of data for the No Adult Talk, Adult to Adult Talk, and Parent Infant Talk circumstances respectively.

### Coding

Coding focused on determining counts for the protophones, cries, whimpers, and laughs, which together accounted for 99% of all utterances. The three precanonical protophone types that were coded for inclusion in the analysis (squeals, growls and vowel-like sounds) were collapsed together. Cries and whimpers (for definition of the distinction, [50]) were also collapsed into a single distress category.

Protophones are spontaneous: no particular emotional state or stimulus is needed to produce them. Thus, they provide a basis upon which speech development depends since it must be possible to produce any element of speech in any emotional state as well as in a state of neutrality or pure interest in self-produced sound. Cry/whimper and laughter, on the other hand, have clear correlates in other species and are associated in infants with unambiguous states of distress (negativity) or exultation/joy (positivity).

Coders were encouraged to work intuitively in differentiating protophones from cry/whimper and laughter, with training criteria and procedures specified in prior studies [23, 45]. As will be seen below, coder agreement on the distinctions is good, in spite of mixtures among types that yield intermediate cases.

Protophones, cry/whimper, and laughter were all counted in accord with a “breath group” criterion [51]: each voiced period produced on a single egress was counted as one utterance. Thus, all three utterance types were counted in a similar way, breaking bouts of cry/whimper or laughter into groupings of roughly similar dimensions to bouts of protophones.

After the 5-minute coding period for each all-day recording segment, coders responded to the following questions (among others not relevant here): 1) Did any other person talk to the baby? This could be the parent or another adult or child. 2) Do you think the baby was alone in the room? And 3) do you think the baby was asleep? The questions were answered on a 5-point scale, where 1 indicated Never, 2 Some of the time, 3 About half the time, 4 Most of the time, and 5 The entire time.

Coders for both all-day and laboratory recordings were 16 normally-hearing female students from the University of Memphis School of Communication Sciences and Disorders, who been trained in phonetic transcription during their program of study. The additional six-to eight-week training for the coding of infant vocalizations is described in detail in prior publications [23, 45]. The set of segments corresponding to recordings from each infant was assigned to a single coder. The protocol specified that coders should work through the entire data set for each infant to which they had been assigned before proceeding with the next infant. Coding of each recording was completed before coding of another recording was begun, and the (21 for Atlanta or 24 for Memphis) five-minute segments were coded, and questionnaires were answered for each segment in the chronological order in which they had occurred during the recording day.

### Coder agreement

There are extensive data on the agreement among the coders in prior publications [23, 45]. Since laughter was not considered in those publications, we amplify the findings of prior agreement analysis here. 523 five-min segments were coded independently by two of the coders, with 12 of the 16 participating, each being semi-randomly assigned to segments from six different ages and at least 4 different infants for the agreement phase of the coding. The correlations between counts for the different coders were: protophones: *r* = .84, *ρ* = .91; cry/whimpers: *r* = .94, *ρ* = .77; laughs: *r* = .89, *ρ* = .67. Restricting the data to the second half-year only, when laughter is much more common than earlier, the 293 five-min segments showed correlations of: protophones: *r* = .84, *ρ* = .92; cry/whimpers: *r* = .84, *ρ* = .77; laughs: *r* = .93, *ρ* = .73.

## Results

Figure 1A shows the relative rates of protophones, cry/whimpers, and laughs in the Atlanta data, with laughter showing rates so low that their divergence from 0 is hard to discern on the graph prior to the middle of the first year. In the second half-year, >1400 laughs occurred during wakeful segments, but cry/whimpers were >8 times more frequent (>12,000), and protophones ∼74 times more frequent (>106,000) than laughs. To place the rates in perspective, laughs occurred on average about 3.8 times per hour in the second half-year, while protophones nearly 292 times per hour.

**Figure 1:**
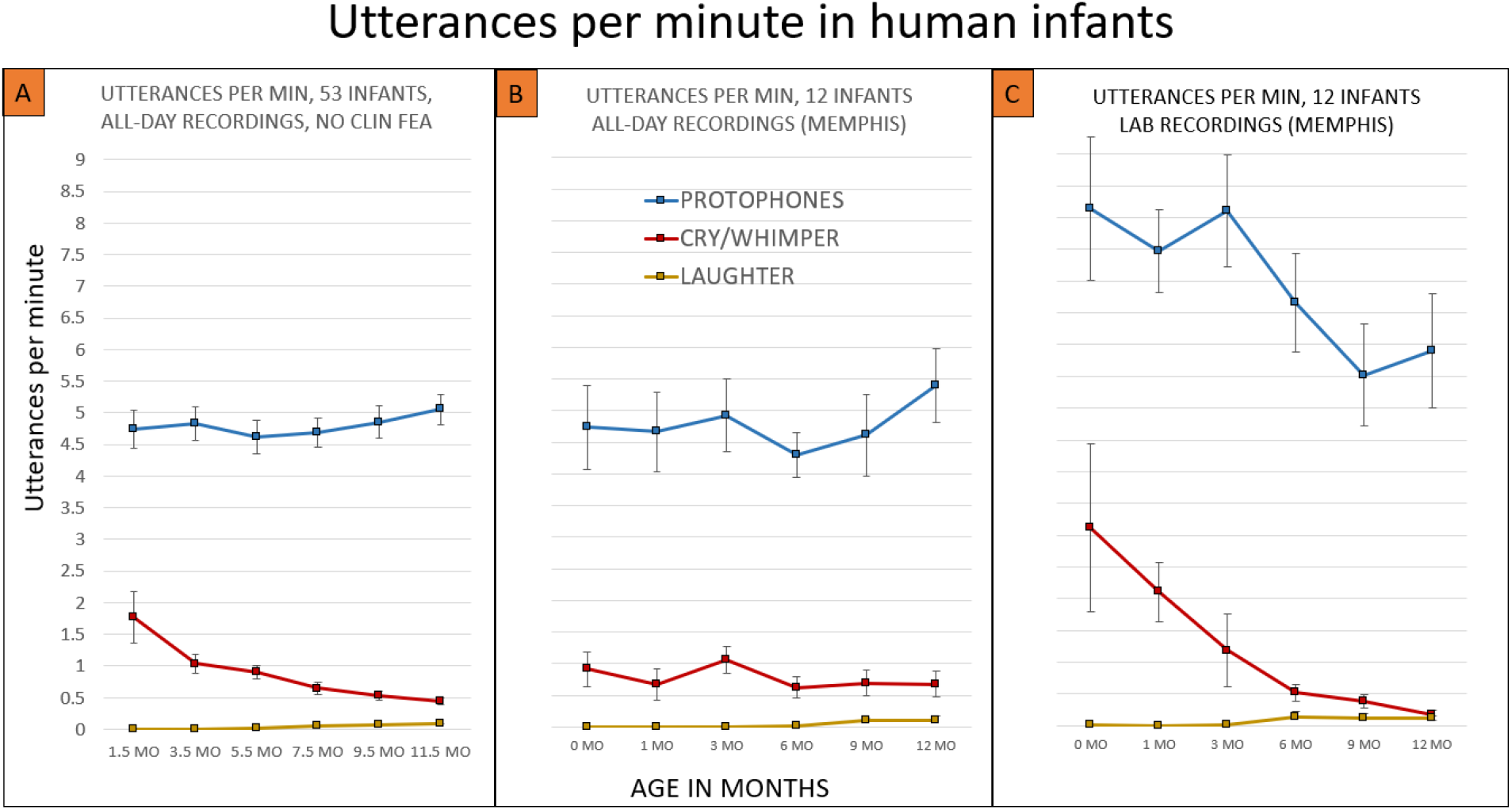
Rates of protophones, cry/whimpers, and laughs in human infants across the first year. A) Based on all-day recordings of 53 infants determined to have no clinical features (to be typically developing) in the Atlanta sample, protophones were massively more frequent than cry/whimpers, and cry/whimpers were massively more frequent than laughs. Standard error bars illustrate the reliability of these differences, although the standard errors were so small for the laughs that they are contained within the square markers at each age. B) Based on all-day recordings of 12 typically-developing infants in the Memphis sample, the patterns were similar to those from the Atlanta sample, although predictably, given the smaller sample size, the error bars are larger. C) Based on laboratory samples for the same 12 infants in Memphis, the patterns confirm those from the all-day recordings, although rates of protophones and cries were lower at the older ages than the younger ones in the laboratory samples. Means and standard errors for Figure 1 were computed at the infant level in each relevant sample (mean across infants and SD divided by the square root of the number of infants).

The Memphis data based on all-day recordings are displayed in Figure 1B, supporting the basic patterns of the larger Atlanta sample. In the second half-year laughs were infrequent (N = 228) compared with protophones (14,658 or 64 times more frequent than laughs) and cry/whimpers (2043 or 9 times more frequent than laughs).

The circumstances of recording played a role in the frequency of occurrence of utterances as illustrated in Figure 1C, where it can be seen that the 12 Memphis infants in the laboratory setting produced more protophones than in the all-day recordings. They also produced more protophones and more cry/whimpers early in the year than later. Yet even in the laboratory setting, laughs were very infrequent compared with the other vocal types. In the second half-year, protophones (13,396) were 50 times more frequent than laughs (N = 268) and cry/whimpers (784) were 2.9 times more frequent than laughs.

The reduction in frequency of occurrence across age for protophones in the laboratory recordings (Figure 1C) can be interpreted, we think, primarily as being a result of the greater mobility of infants, who in the second half-year tended to crawl or walk about the playroom finding toys and other objects to explore—mothers in the second half-year seemingly had less influence on the infants’ vocalizations except in the Parent Infant Talk circumstance.

Figure 2A illustrates a salient effect of parent-infant interaction, where laughter was indeed far more frequent in the second-half year during the Parent Infant Talk sessions than during the sessions with little or no IDS. The existence of even a small amount of laughter in the other in the No Adult Talk and Adult to Adult Talk sessions may be due to the fact that parents occasionally violated recording protocol and spoke to infants briefly. Notice that the scales are different for Figure 2A vs. 2B and 2C: Regardless of circumstances or age, protophones (Figure 2C) were >14 times more frequent than laughs in the laboratory at every age and every circumstance. The very high rate of cry/whimper at the youngest age (Figure 2B) in the No Adult Talk circumstance can be attributed, we think, to infant protest at being left in a crib or stroller with little or no adult attention—mothers did not allow the crying to go on too long, choosing to hold the infant while reading if the infant persisted in crying. Figure 2A shows that at the latest ages, laughter proved to be competitive with cry/whimper for rate of vocalization in the Parent Infant Talk circumstance.

**Figure 2:**
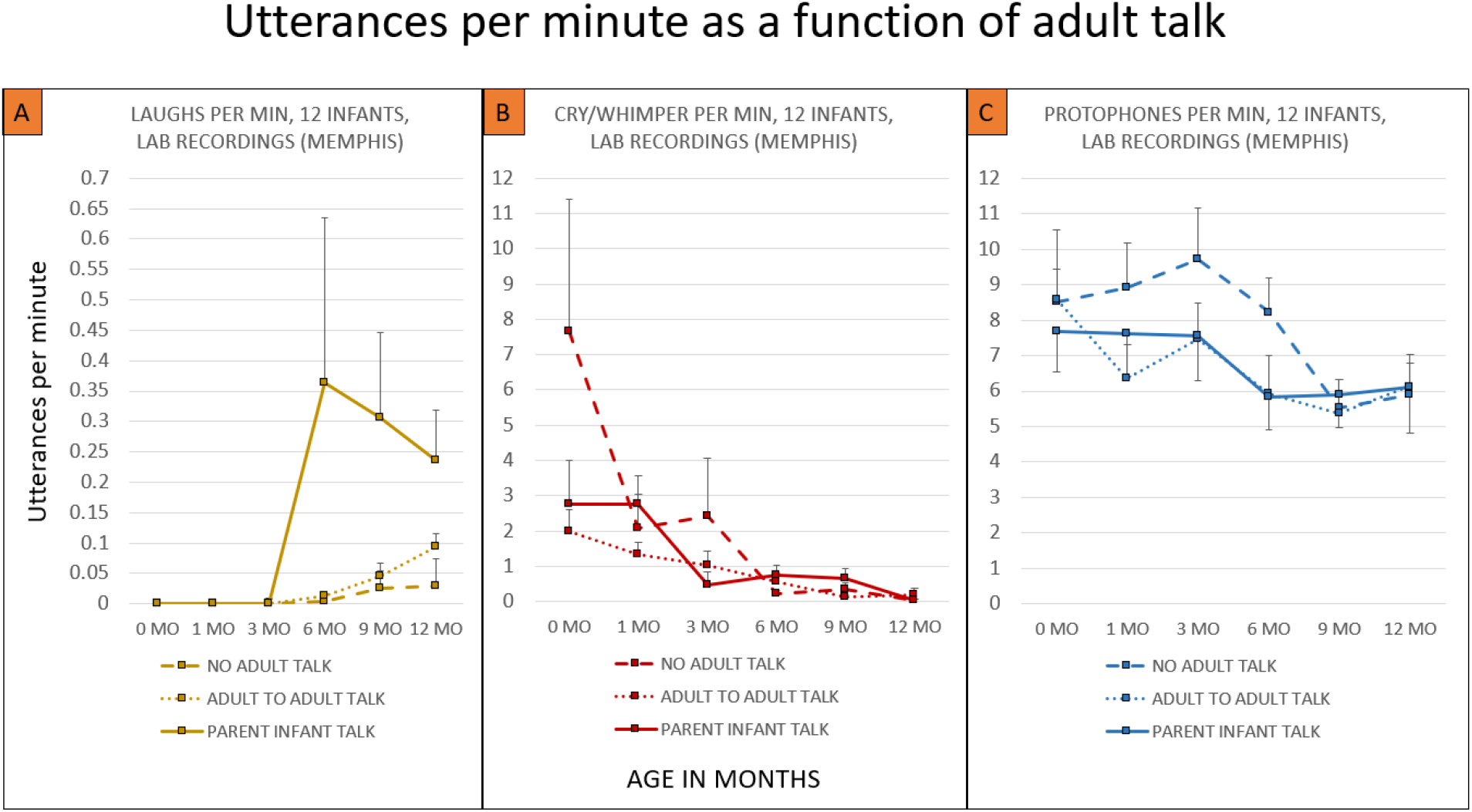
Rates of the three vocal types based on three different laboratory recording protocols. A) Laughs occurred infrequently in all three protocols in the first three months, but although the frequency was very low compared with cry/whimpers and protophones in the second half-year (note the y-axis scale differences for Figure segment A vs. Figure segments B and C), the rates of laughter were, as expected, at their highest in the Parent Infant Talk circumstance (infant-directed speech). B) Cry/whimpers were far more frequent than laughs through 3 mo, but the rates were more comparable in the second half-year. Cry/whimper rates based on the laboratory counts should, however, be interpreted in light of the fact that the recordings were sometimes interrupted to soothe a crying or fussing infant. The high rate of cry/whimper during the No Adult Talk circumstance at 0 months appears to have been the result of infant distress at being left nearby but unattended in the room, which resulted in either the parent deciding to hold the infant during No Adult Talk or interruption of the recording to calm the infant. C) Protophone rates were higher in the laboratory than in the all-day recordings though they tended to fall across the first year. Note that Parent Infant Talk did not in general correspond to notably higher rates of protophones than in the other circumstances, a fact we interpret as corresponding to the largely endogenous nature of protophone production. Means and standard errors for Figure 2 were computed at the infant level.

Finally, Figure 3 shows data based on the rates of laughter occurring in the all-day recordings as a function of the amount of infant-directed-speech (IDS) and whether the infant was alone in the room during the five-minute segments as indicated by the questionnaire items administered during coding. The results support the recognition that laughter is a social phenomenon, with several times more laughter having occurred with IDS and when infants were not alone than otherwise. At the same time, even in five-minute segments where some one was talking to an infant, the rate of laughter was miniscule compared with protophone rates.

**Figure 3:**
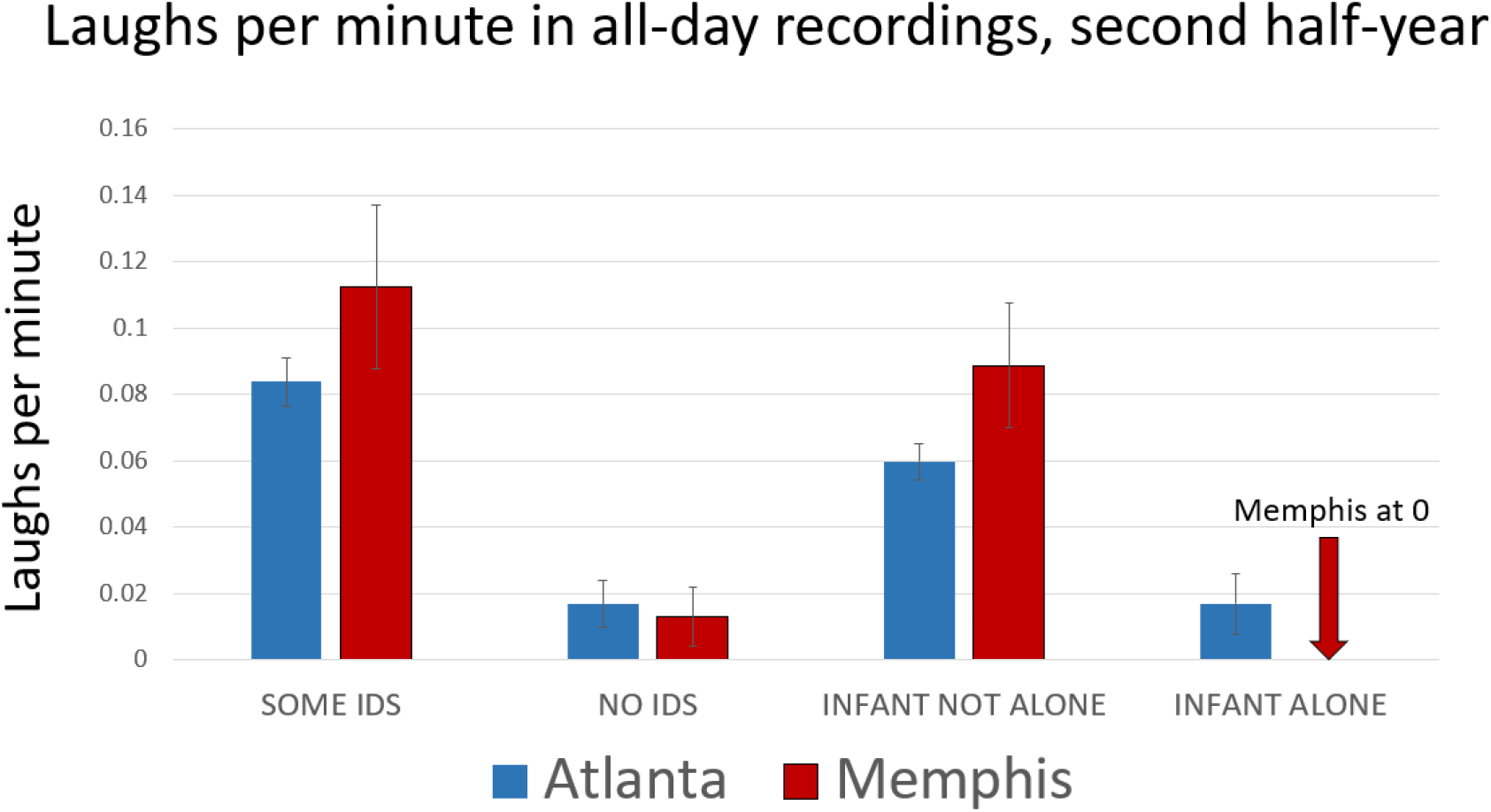
Rates of laughter during all-day recordings in two circumstances in the second half-year. The coding allowed segregation of the data according to whether there was infant-directed speech (IDS) or not and also according to whether the infant was deemed alone in the room or not. The comparisons from both Atlanta and Memphis data are consistent with the social nature of infant laughter, which occurred more frequently with IDS and during periods when the infant was awake but not alone. Still there was very large variation among the infants and across segments where laughter occurred. More than 90% of segments even in the second half-year contained no laughter at all, while occasional segments (∼1%) included at least 10. By contrast, more than 90% of segments included at least one protophone and ∼2/3 included more than 10. Means and standard errors for Figure 2 were computed at the segment level.

## Discussion

### The infrequency of laughter

Our report is not intended to deemphasize the importance of laughter in human infant development. The salience of laughter events that sometimes occur repeatedly in playful interactions between parents and infants (in peekaboo, for example) provide intuitively persuasive evidence that bonding and social learning may be richly served by such joyful interactions. Yet the infrequency of laughter occurrences based on this extensive sampling from all-day recordings, a rate at least 50 times lower than that of protophones, is surely surprising. It is important to note that the rates of laughter reported here for laboratory recordings are *not* low compared with rates that have been reported previously based on experimental observations of parent-infant interaction. In fact, the rates in the second half-year for the Parent Infant Talk circumstance in the Memphis data appear to be a little higher than those reported in the most comprehensive previous study we know of reporting infant laughter rates [41]. In the presence of a mother not engaged in IDS, the study reported laughter rates lower than in the present data.

Infant laughter is salient not only in humans but also in other apes [52]. In the only direct quantitative comparison we know of across human and non-human ape infants [24], we found that 3 bonobo infants laughed during rough and tumble play or tickling and that laughter appeared to be the most frequent type of vocalization in the bonobo infants. The sample size was insufficient to make useful statistical comparisons of rates of laughter in the human and bonobo infants, but protophones in the humans were far more frequent than laughter in either case. Playful laughter has been reported for all the great apes and for many mammals [52, 53], and has been speculated to provide a phylogenetic platform for evolution of language [39, 40, 54]. Yet its occurrence in human infants was shown here to be remarkably rare especially when compared with the protophones, the precursors to speech.

### The high frequency of protophones and the endogenous nature of vocal development

The massive rate of protophone production along with the fact that protophones are overwhelmingly directed to nobody during playful exploration from the beginning of human life and through the first year compel us to recognize that the activity is predominantly generated endogenously and in comfort. Laughter and cry/whimper, on the other hand, are both generated primarily in situations of either social play or of distress. These more emotionally grounded signals play the same kind of role in humans that similar vocalizations play in other mammals, and prominently in the great apes. But protophones are at best minimally present in other apes [24], and to the extent that they may occur, they have never been shown to exhibit the exploratory characteristic that is in fact the predominant mode of production of protophones in the human infant.

We have long argued that the ability and inclination to produce protophones copiously forms a foundation without which the development of language would be impossible [55, 56]. The reason is simple and logical: Language elements are always possible to produce in any circumstance of emotion or illocutionary intent—the word “elephant” can be produced to complain, to request, to name, to correct, to criticize, to teach, to practice the pronunciation of the word, and in any state of pleasure or displeasure. If it were not so, “elephant” would not be a word and could not pertain to the lexicon of any language. Thus, the ability to produce a set of particular sounds freely in any emotional state is clearly a foundation without which learning to use a word would be impossible. We call this capability to produce particular sounds in any emotional state “vocal functional flexibility” (VFF), and have proven it to be present extensively in the human infant’s protophones in the first months of life [21, 57]. Laughs and cry/whimpers in infancy do not show VFF.

### Fitness signaling and the origin of language

Why, then, do protophones exist at all? And especially why do they occur so frequently compared with important vocalizations such as laughter? The questions are not trivial because it can be assumed that the ability to produce sounds with VFF must have preceded the origin of vocal language, which as noted, is logically dependent on VFF. Consequently, at their earliest appearance in hominin evolution, vocalizations with VFF must have been selected for in accord with pressures that had nothing to do with language, which did not yet exist.

The evolutionary origins of laughs and cry/whimpers, in contrast, fit the more standard mold of presumable selection pressures. Both these types of vocalizations express definable emotional states and serve definable and relatively consistent functions that have direct potential benefits at the time they are produced. Cry/whimpers signal need for care, and laughter (especially since it seems to be elicited specifically in social circumstances) signals social connection. It seems straightforward to postulate that any mammal, being dependent on maternal care, is under selection pressure to have the ability to respond with these kinds of sounds as needed in appropriate situations.

But the protophones are different because they do not have a fixed valence or a predominant immediate social function that could have been the basis for selection. Even a comfortable infant who is entirely alone produces massive numbers of protophones, and even if parents are talking to an infant, most of the infant’s protophones are not directed to anyone [25]. So, we reason, the predominant function of the protophones must be based on advantages that do not tend to accrue in the immediate context of their production. Rather, we argue, the protophones predominantly supply information about infant wellness even to caregivers who are busy doing something else nearby.

This kind of fitness signaling has been argued to be particularly important to human infants for two reasons. First, human infants and their ancient hominin precursors are and were more altricial than other apes, with much longer developmental periods of helplessness and need for provisioning by others than in the case of other apes [58]. Thus, pressure on signaling their wellness may have resulted in the ancient hominin infant vocal system being selected for engagement with the same motivational/emotional system that generates exploration with the hands in other baby primates. We presume that sounds produced by the infant hominin’s own phonatory system came thus to be objects of exploration and play [59, 60]. The capability and inclination to produce these sounds, and thus to indicate their wellness, presumably put them at an advantage with respect to other hominin infants in the competition for investment by provisioning from others and in the competition to be kept rather than abandoned in times of stress.

A second reason that the pressure on such fitness signaling may have been particularly high in hominin infants is that ancient hominin groups were larger than those of other apes, and increasingly so over the evolution of the hominins [61]. These larger groups were also presumably increasingly cooperative breeders, which is to say infants were cared for and provisioned not just by the mothers, but by alloparents of the group, a pattern of rearing that is seen strongly in just one other group of primates, the New world callitrichids. Importantly, the callitrichids are also the only other group of primates known to engage in “babbling” in infancy [62, 63]. We reason, along with others, that the pressure on vocal fitness signaling is deep in the hominin line both because of altriciality and because of cooperative breeding, given that infants could profit from broadcasting fitness indicators in the competition for care from a variety of alloparents [64].

Of course there are many fitness indicators: the color of the skin, the vigor of movement, the ability to raise the head, to move the fingers, and so on. All these presumably play roles in how caregivers of various mammalian species determine their investments in the young who are in their charge. The protophones offer a special leg up on fitness signaling, however, because they can occur even when the potential caregiver is not attending to them, for example, after putting the infant down to forage. We reason that the value of these vocal signals may be recognized even if semiconsciously, accumulating in the awareness of the potential caregiver, who may provide benefit to the infant much later.

Our proposal emphatically does not suggest that the protophones constitute language. Rather, we propose that the ability and inclination to produce protophones supplies a platform on which later development can build. Further, ancient hominin infants, according to our proposal, were selected to produce protophone-like sounds first and later came under additional natural selection pressures for more elaborate communication. Vocal language would not be possible without a foundation of functionally flexible vocalization, but much remains to be evolved and developed beyond the achievement manifest in the protophones.

